# Extant interspecific hybridization among trematodes within the *Schistosoma haematobium* species complex in Nigeria

**DOI:** 10.1101/2023.07.24.550253

**Authors:** OG Ajakaye, EE Enabulele, JB Balogun, OT Oyeyemi, ME Grigg

## Abstract

**Background:** Natural interspecific hybridization between the human parasite (*Schistosoma haematobium* [Sh]) and bovine parasites (*S. bovis* [Sb]*, S. curassoni* [Sc]) is increasingly reported in Africa. We developed a multi-locus PCR DNA-Seq strategy that amplifies two unlinked nuclear (*transITS*, *BF*) and two linked organellar genome markers (*CO1*, *NAD5*) to genotype *S. haematobium* eggs collected from infected people in Ile Oluji/Oke Igbo, Ondo State (an agrarian community) and Kachi, Jigawa State (a pastoral community) in Southwestern and Northern Nigeria, respectively.

**Principal Findings:** We applied this methodology against 57 isolates collected from a total of 219 participants. All patients from Jigawa state were infected with just one of two haplotypes of an *S. haematobium x S. bovis* hybrid based on sequences obtained at *CO1*, *NAD5, transITS* and *BF* markers. Whereas samples collected from Ondo state were varied. Mitonuclear discordance was observed in all 17 patients, worms possessed an *Sb* mitochondrial genome but one of four different haplotypes at the nuclear markers, either admixed (heterozygous between *Sh x Sc* or *Sh x Sb*) at both markers (n=10), *Sh* at *BF* and admixed at *transITS* (*Sh x Sc*) (n=5), admixed (*Sh x Sc*) at *BF* and homozygous *Sc* at *transITS* (n=1) or homozygous *Sh* at *BF* and homozygous *Sc* at *transITS* (n=1).

**Significance:** Previous work suggested that zoonotic transmission of *S. bovis* in pastoral communities, where humans and animals share a common water source, is a driving factor facilitating interspecific hybridization. However, our data showed that all isolates were hybrids, with greater diversity identified in Southwestern Nigeria, a non-pastoral site. Further, one patient possessed an *S. bovis* mitochondrial genome but was homozygous for *S. haematobium* at *BF* and homozygous for *S. curassoni* at *transITS* supporting at least two separate backcrosses in its origin, suggesting that interspecific hybridization may be an ongoing process.

**Author Summary:** Interspecific hybridization between trematode parasites poses serious health risks to humans. Many systems have shown possible hybridization between different schistosome species. As evidence of natural hybridization between human *S. haematobium* and animal *S. bovis* or *S. curassoni* has grown in recent years, epidemiological surveys across potential hybrid zones are required, particularly in endemic African regions. According to several reports, indiscriminate human-animal water contact is a major factor contributing to hybridization of human and animal schistosomes. We collected and genotyped 57 parasite isolates from pastoral and non-pastoral communities in Kachi, Jigawa state, and Ile Oluji/Oke Igbo, Ondo state, Nigeria to screen for hybrids. In both sites, we found *Schistosoma* hybrids with mitonuclear discordance and repeated backcrossing between *S. haematobium*, *S. bovis*, and *S. curassoni*. Contrary to previous reports, *Schistosoma* hybrids appear to be widespread and not solely dependent on human-animal water interactions.

## Introduction

Schistosomiasis is a highly prevalent water transmitted disease classified by the World Health Organization (WHO) as neglected. It is the second most prevalent tropical disease caused by five major species of flat worms, specifically *Schistosoma mansoni, S. haematobium, S. japonicum, S. mekongi,* and *S. intercalatum*. The first two species cause approximately 20 million infections in Africa and are associated with severe chronic health consequences in affected populations [1&2]. Many other schistosome species, including *S.bovis*, *S. curassoni, S. mattheei* are known to infect livestock especially in sub–Saharan Africa [3]. Nigeria has the highest number of schistosomiasis cases in the world, with over 26% of Nigerians requiring chemotherapy. Despite annual mass chemotherapy programs, the urogenital form of schistosomiasis, caused by *S. haematobium*, remains a significant public health problem, with a current national prevalence of about 10% [4].

In 2009, field evidence of hybridization between *S. haematobium* and the bovine schistosome *S. bovis* was reported for the first time in Africa based on typing strategies using the mitochondrial marker *Cox1* (CO1) and the *transITS* (ITS) nuclear marker, located within the ribosomal RNA gene array [5]. Since the 2009 report, hybrid *S. haematobium* infections are now widely reported in countries in Africa, including Nigeria [6, 7,]. Whether such interspecific hybridization impacts urogenital schistosomiasis transmission, virulence, and treatment drug efficacy remains enigmatic

More recently, several genome sequencing projects investigated *S. haematobium* [8–10] and *S. bovis* [11] at whole genome resolution to infer genome ancestry. Hybrid ancestry was extant, but the proportion of the nuclear genome derived from *S. bovis* was limited, resulting in only 8.2% [8] or 23% [10] within the *S. haematobium* genomes investigated. The failure to identify full genome hybrids was interpreted to suggest that interspecies hybridization between *S. haematobium* x *S. bovis* is ancient, that it has occurred only rarely, and that *S. bovis* derived genome blocks have been purified by ongoing hybridization with *S. haematobium,* the more relevant schistosome species naturally infecting humans. It was also hypothesized that hybridization was endemic and occurring where the cultural practices of humans and livestock sharing the same water bodies is common [5, 12–14].

In Nigeria, nomadic farming and pastoralism are a common cultural practice, particularly in the northern regions of the country. Pastoral animals are grazed around rivers and streams which also serve as the main sources of water for the communities. Therefore, these regions exist as potential high-risk areas for hybridization between human and livestock schistosome species. To test this hypothesis, that interspecific *S. haematobium* hybrids possessing a mixed ancestry with bovine schistosomes are common in water bodies shared between humans and cattle, we sampled two ecologically distinct *S. haematobium* endemic communities in Southwestern and Northern Nigeria. The sampled community in southern Nigeria is devoid of livestock whereas the community in the northern part of the country is pastoral. We report here the genetic profiles of the *S. haematobium* from the two locations using genetic markers anchored in the mitochondria (*CO1*, *NAD5*) and two unlinked nuclear genome markers (trans*ITS*, *BF*). The identification of homozygous alleles derived from animal schistosome species in sites devoid of livestock supports the notion that ongoing hybridization and backcrosses with animal schistosome species is occurring, independent of human-animal freshwater usage. Our work highlights the need to apply modern high throughput techniques to screen for hybrids to assess the degree to which ongoing hybridization is occurring between animal and human schistosomes, and to identify the relevant hosts and infection reservoirs that are promoting these interactions.

## Results and Discussion

### Field Site data

Interspecific hybridization between animal and human schistosomes is thought to occur preferentially in pastoral sites where animals and people share water bodies. To assess the degree to which this was occurring in Nigeria, two ecologically distinct *S. haematobium* endemic communities in southwestern and northern Nigeria were sampled (Figure 1). In the community in southwest Nigeria that was devoid of livestock, 17 of 107 filters contained parasite eggs. DNA was extracted from these *S. haematobium* positive urine samples. Forty of 112 filters contained parasite eggs in the pastoral community in northern Nigeria, and DNA was extracted from these *S. haematobium* positive urine samples. In aggregate, the prevalence of urogenital schistosomiasis infection was higher at the pastoral site in Kachi (35.7%) compared to the agrarian site in Ile Oluji/Oke Igbo (15.9%). The age of the participants ranged from 3-19 years, prevalence was higher in males (55.8%) than females (45.2%), but this did not reach significance (Table 1). Age was significantly associated with infection (p<0.00001), as has been reported previously. In both study sites, a heavy intensity of infection was recorded (≥50 eggs per 10 ml of urine) (Table 1).

**Figure 1.**
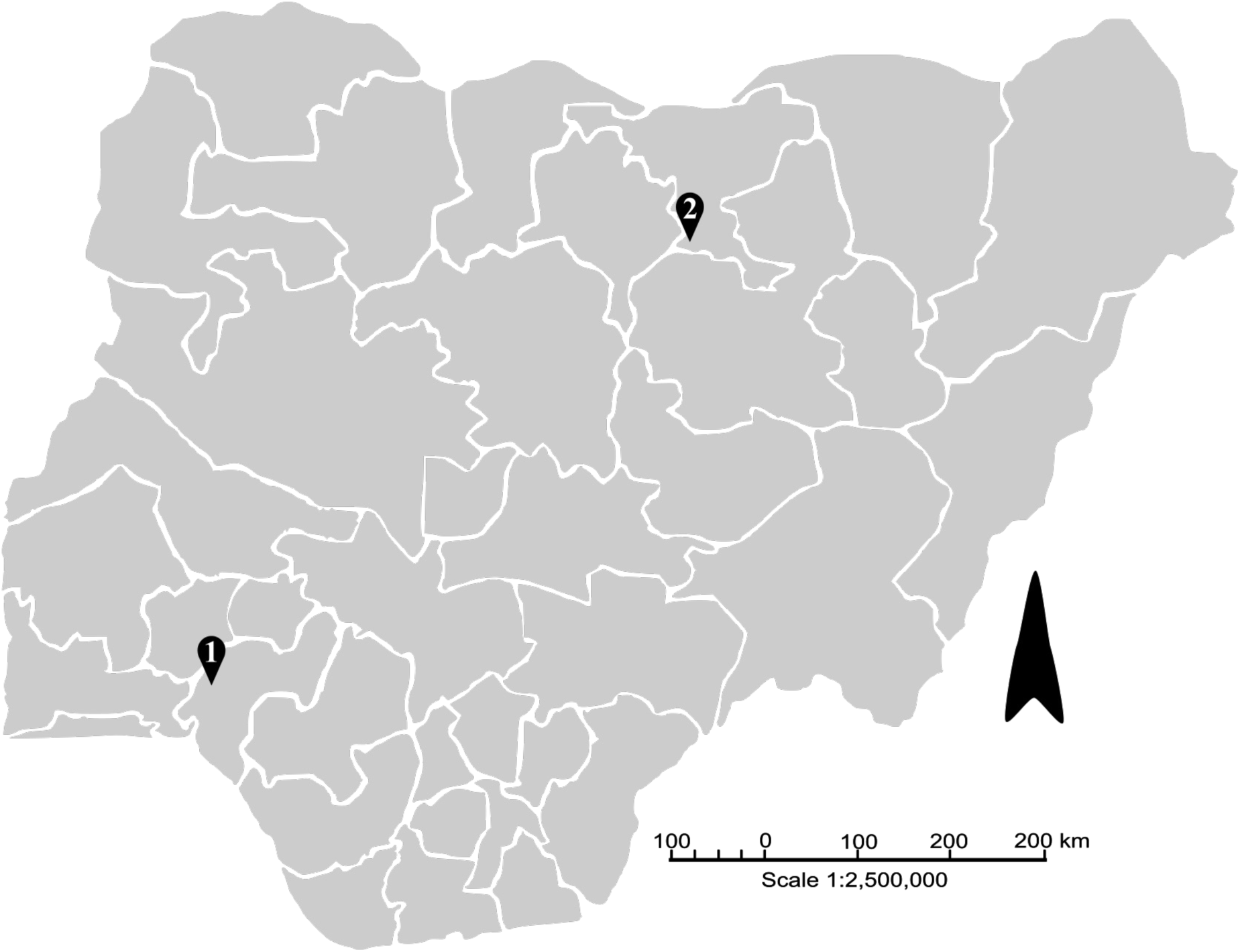
Map of Nigeria showing the 36 states and the specific sampling locations in Ondo State (1), Ile Oluji/Oke Igbo [latitude 5° 45N and 8° 15N and longitude 4° 30E], and Jigawa State (2), Kachi [latitude 11° 73’ N and longitude 9° 33’ E] for *S. haematobium*. Arrow depicts North.

**Table 1:** *Schistosoma haematobium* prevalence in Ile Oluji/Oke Igbo (Ondo State) and Kachi (Jigawa State), Nigeria.

| Variable | Category | Locations |  | Total | P value |
| --- | --- | --- | --- | --- | --- |
|  |  | Ile Oluji/Oke Igbo | Kachi |  |  |
| Human Infection | Negative | 90 (84.1) | 72 (64.3) | 162 (74.0) | 0.0008* |
|  | Positive | 17 (15.9) | 40 (35.7) | 57 (26.0) |  |
| Infection Intensity | Heavy | 11 (64.7) | 17 (42.5) | 28 (49.1) | 0.1249 |
|  | Light | 6 (35.5) | 23 (57.5) | 29 (50.9) |  |
| Gender | Female | 54 (50.5) | 45 (40.2) | 99 (45.2) | 0.1262 |
|  | Male | 53 (49.5) | 67 (59.8) | 120 (55.8) |  |
| Age | < 10 | 73 (68.2) | 22 (19.6) | 95 (43.4) | <0.00001* |
|  | > 10 | 34 (31.8) | 90 (80.4) | 124 (56.6) |  |

\* $P < 0.05$ , Prevalence percentages are in parenthesis

### Schistosoma haematobium genotyping

To investigate whether interspecific schistosome hybrids were causing human infection at the two study sites, a PCR-DNA sequencing strategy was employed to determine the alleles present within the nuclear genome marker *transITS* and the mitochondrial genome marker *CO1.* In Kachi, all patients were infected by interspecific hybrids, the result of a cross between *S. bovis* x *S. haematobium* based on the two sequences obtained at the *transITS* marker on chromosome 2 (Table 2). Of particular interest, a private SNP at nucleotide 502 (based on GenBank reference sequence MH014047) was identified that was unique to strains circulating in Kachi that has not been previously observed in any published *S. haematobium* or *S. bovis* allele (Suppl.Table 2). It was not possible to interpret the origin of the private SNP, *i.e.,* whether it was of *S. bovis* or *S. haematobium* ancestry. At *CO1*, all isolates had an *S. haematobium* haplogroup I allele (Figure 2). One isolate possessed a single, unique (or private) SNP at *CO1* indicating that more than one haplotype was circulating among the patient samples (Figure 3).

**Figure 2.**
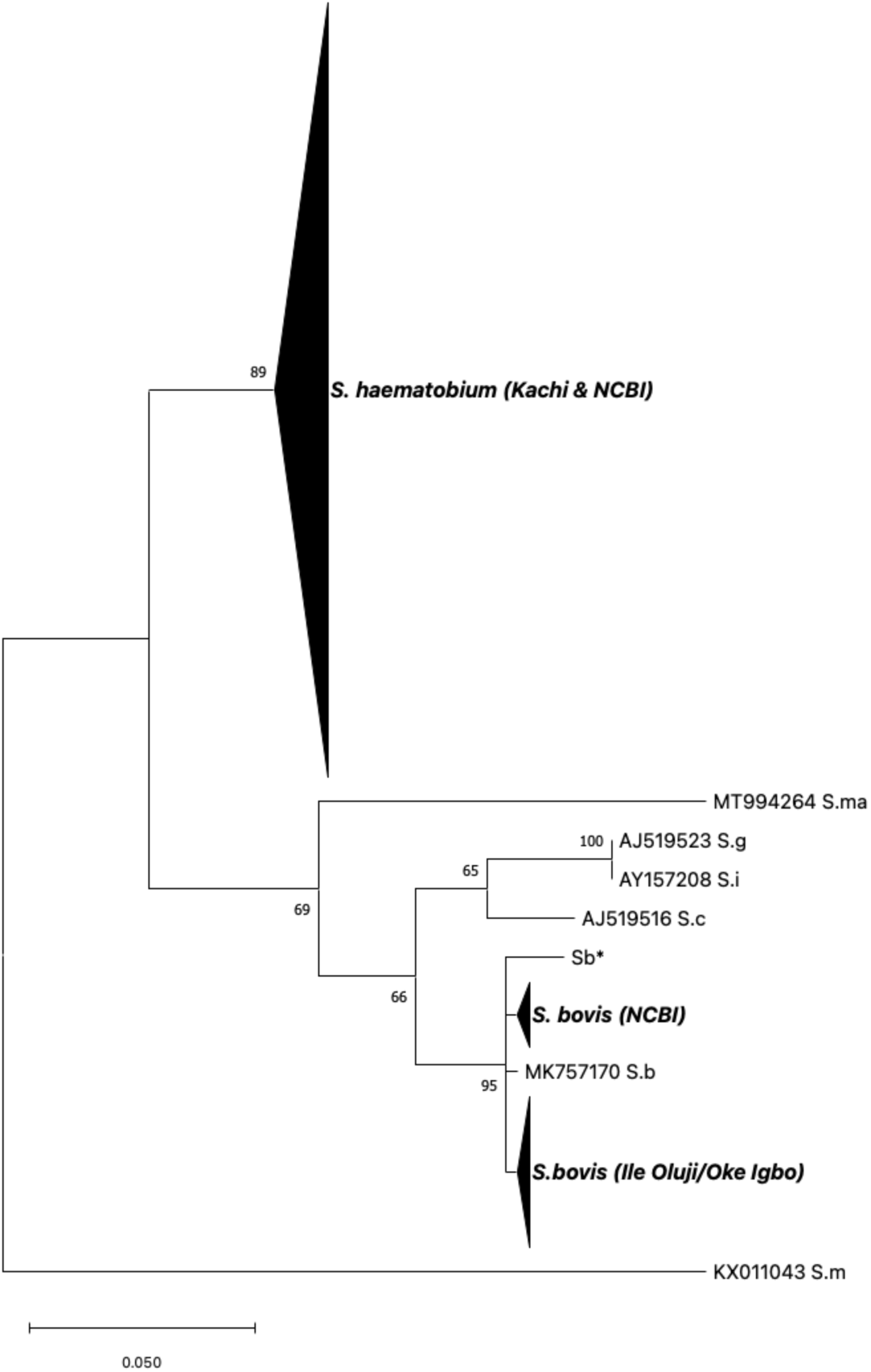
Maximum likelihood tree based on CO1 sequences obtained from *S. haematobium* isolates collected in Oluji/Oke Igbo and Kachi in Nigeria. Reference sequences from worms within the *S. haematobium* species complex were added for comparative purposes, and are comprised of published sequences retrieved from GenBank. The OC1 sequence from *S. mansoni* (KX011043) was used as the outgroup. (Sb* - this study, *S. bovis* worm from cow recovered from an abattoir in Nigeria); *S.b* = *S. bovis; S.m* = *S. mansoni*; *S.c* = *S. curassoni*; *S.ma* = *S. mattheei*; *S.g* = *S. guineensis*; *S.i* = *S. intercalatum*.

**Figure 3.**
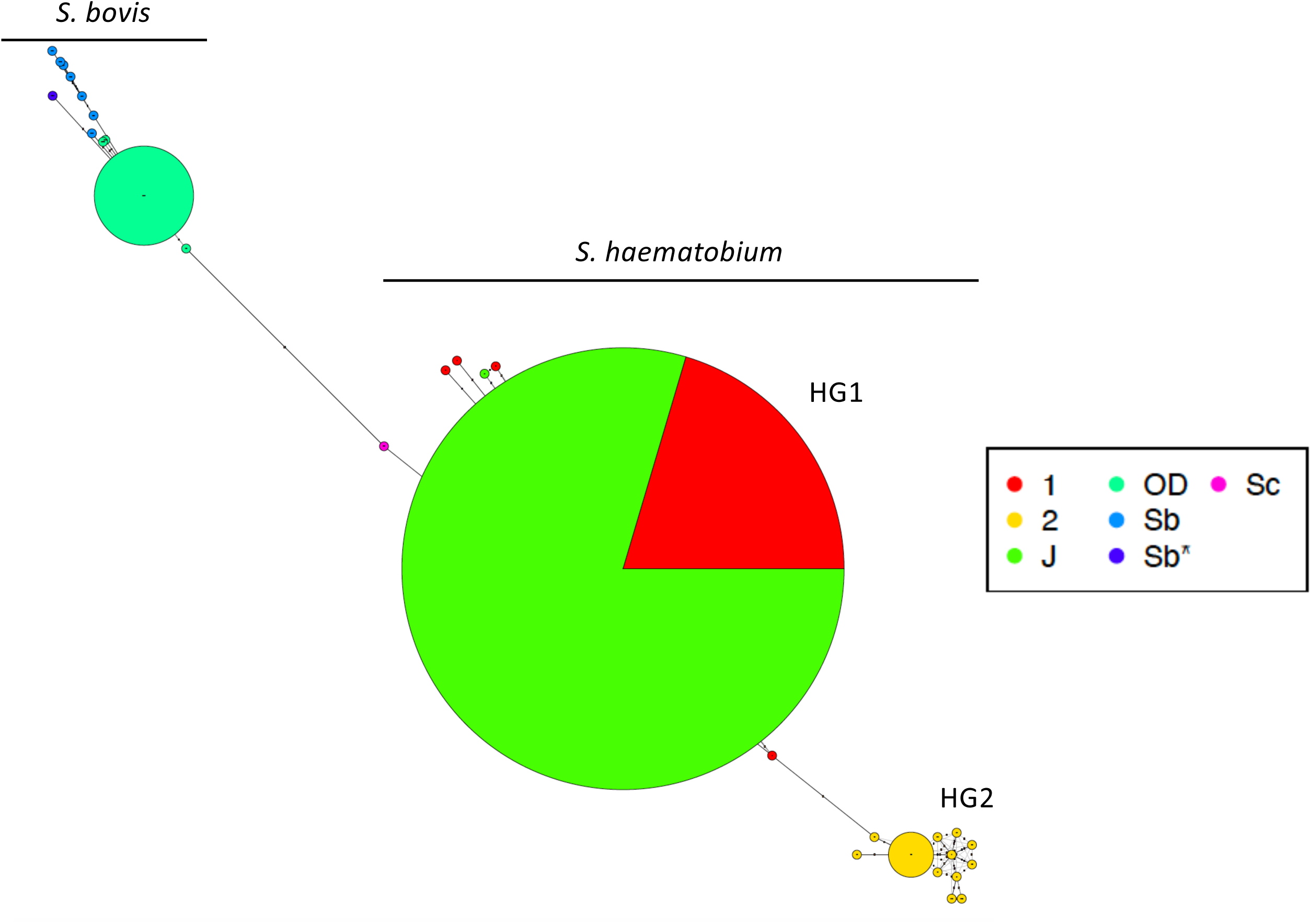
Haplotype network at *CO1*. *S. haematobium CO1* alleles circulating among isolates recovered from patients in Kachi, Jigawa state (J) belonged to the *S. haematobium* group 1 (HG1) haplotype (green). Red – *S. haematobium* haplogroup 1 reference sequences obtained from GenBank. For reference, *S. haematobium* group 2 (HG2) haplotypes (yellow) are also depicted for sequences obtained from GenBank. *S. bovis CO1* alleles circulating among isolates recovered from patients in Oluji/Oke Igbo, Ondo state (OD) were distinct (seagreen) from sequences obtained from GenBank annotated as *S. bovis*. Sb* represents the allele identified at *CO1* for the *S. bovis* isolate recovered from an abattoir in Nigeria, that served as a control sequence for *S. bovis*.

**Table 2:**

| Isolate | Location | State | Source | Nuclear Markers |  | Mitochondrial Markers |  |
| --- | --- | --- | --- | --- | --- | --- | --- |
|  |  |  |  | *transITS* | *BF* | *CO1* | *ND5* |
| OD1 | Ile Oluji/Oke | Ondo | Human | *S.h* x *S.c* | *S.h* | *S.b* | *S.b* |
| OD2 | Ile Oluji/Oke | Ondo | Human | *S.h* x *S.c* | *S.h* x *S.b* | *S.b* | *S.b* * |
| OD3 | Ile Oluji/Oke | Ondo | Human | *S.h* x *S.c* | *S.h* | *n.d.* | *S.b* |
| OD4 | Ile Oluji/Oke | Ondo | Human | *S.h* x *S.c* | *S.h* | *S.b* | *S.b* |
| OD5 | Ile Oluji/Oke | Ondo | Human | *S.h* x *S.c* | *S.h* | *S.b* | *S.b* |
| OD6 | Ile Oluji/Oke | Ondo | Human | *S.h* x *S.c* | *S.h* x *S.b* | *n.d.* | *S.b* |
| OD7 | Ile Oluji/Oke | Ondo | Human | *S.h* x *S.c* | *S.h* | *S.b* | *S.b* |
| OD8 | Ile Oluji/Oke | Ondo | Human | *S.h* x *S.c* | *S.h* x *S.b* | *S.b* | *S.b* |
| OD9 | Ile Oluji/Oke | Ondo | Human | *S.h* x *S.c* | *S.h* x *S.b* | *S.b* | *S.b* |
| OD10 | Ile Oluji/Oke | Ondo | Human | *S.h* x *S.c* | *S.h* x *S.b* | *S.b* | *S.b* |
| OD11 | Ile Oluji/Oke | Ondo | Human | *S.h* x *S.c* | *S.h* x *S.b* | *S.b* | *S.b* |
| OD12 | Ile Oluji/Oke | Ondo | Human | *S.h* x *S.c* | *S.h* x *S.b* | *S.b* | *S.b* |
| OD13 | Ile Oluji/Oke | Ondo | Human | *S.h* x *S.c* | *S.h* x *S.b* | *S.b* | *S.b* |
| OD14 | Ile Oluji/Oke | Ondo | Human | *S.c* | *S.h* x *S.c* | *S.b* | *S.b* |
| OD15 | Ile Oluji/Oke | Ondo | Human | *S.h* x *S.c* | *S.h* x *S.b* | *n.d.* | *S.b* |
| OD16 | Ile Oluji/Oke | Ondo | Human | *S.h* x *S.c* | *S.h* x *S.b* | *S.b* | *S.b* |
| OD17 | Ile Oluji/Oke | Ondo | Human | *S.c* | *S.h* | *S.b* | *S.b* |
| J1 | Kachi | Jigawa | Human | *S.h* x *S.b* * | *S.h* | *S.h* | *S.h* |
| J2 | Kachi | Jigawa | Human | *S.h* x *S.b* * | *S.h* | *S.h* | *S.h* |
| J3 | Kachi | Jigawa | Human | *S.h* x *S.b* * | *S.h* | *S.h* | *S.h* |
| J4 | Kachi | Jigawa | Human | *S.h* x *S.b* * | *S.h* | *S.h* | *S.h* |
| J5 | Kachi | Jigawa | Human | *S.h* x *S.b* * | *S.h* | *S.h* | *S.h* |
| J6 | Kachi | Jigawa | Human | *S.h* x *S.b* * | *S.h* | *S.h* | *S.h* |
| J7 | Kachi | Jigawa | Human | *S.h* x *S.b* * | *S.h* | *S.h* | *S.h* |
| J8 | Kachi | Jigawa | Human | *S.h* x *S.b* * | *S.h* | *S.h* | *S.h* |
| J9 | Kachi | Jigawa | Human | *S.h* x *S.b* * | *S.h* | *S.h* | *S.h* |
| J10 | Kachi | Jigawa | Human | *S.h* x *S.b* * | *S.h* | *S.h* | *S.h* |
| J11 | Kachi | Jigawa | Human | *S.h* x *S.b* * | *S.h* | *S.h* | *S.h* |
| J12 | Kachi | Jigawa | Human | *S.h* x *S.b* * | *S.h* | *S.h* | *S.h* |
| J13 | Kachi | Jigawa | Human | *S.h* x *S.b* * | *S.h* | *S.h* | *S.h* |
| J14 | Kachi | Jigawa | Human | *S.h* x *S.b* * | *S.h* | *S.h* | *S.h* |
| J15 | Kachi | Jigawa | Human | *S.h* x *S.b* * | *S.h* x *S.b* | *S.h* | *S.h* |
| J16 | Kachi | Jigawa | Human | *S.h* x *S.b* * | *S.h* | *S.h* | *S.h* |
| J17 | Kachi | Jigawa | Human | *S.h* x *S.b* * | *S.h* | *S.h* | *S.h* |
| J18 | Kachi | Jigawa | Human | *S.h* x *S.b* * | *S.h* | *S.h* | *S.h* |
| J19 | Kachi | Jigawa | Human | *S.h* x *S.b* * | *S.h* | *S.h* | *S.h* |
| J20 | Kachi | Jigawa | Human | *S.h* x *S.b* * | *S.h* | *S.h* | *S.h* |
| J21 | Kachi | Jigawa | Human | *S.h* x *S.b* * | *S.h* | *S.h* | *S.h* |
| J22 | Kachi | Jigawa | Human | *S.h* x *S.b* * | *S.h* | *S.h* | *S.h* |
| J23 | Kachi | Jigawa | Human | *S.h* x *S.b* * | *S.h* | *S.h* | *S.h* |
| J24 | Kachi | Jigawa | Human | *S.h* x *S.b* * | *S.h* | *S.h* | *S.h* |
| J25 | Kachi | Jigawa | Human | *S.h* x *S.b* * | *S.h* | *S.h* | *S.h* |
| J26 | Kachi | Jigawa | Human | *S.h* x *S.b* * | *S.h* | *S.h* | *S.h* |
| J27 | Kachi | Jigawa | Human | *S.h* x *S.b* * | *S.h* | *S.h* | *S.h* |
| J28 | Kachi | Jigawa | Human | *S.h* x *S.b* * | *S.h* | *S.h* | *S.h* |
| J29 | Kachi | Jigawa | Human | *S.h* x *S.b* * | *S.h* | *S.h* | *S.h* |
| J30 | Kachi | Jigawa | Human | *S.h* x *S.b* * | *S.h* | *S.h* | *S.h* |
| J31 | Kachi | Jigawa | Human | *S.h* x *S.b* * | *S.h* | *S.h* | *S.h* |
| J32 | Kachi | Jigawa | Human | *S.h* x *S.b* * | *S.h* | *S.h* | *S.h* * |
| J33 | Kachi | Jigawa | Human | *S.h* x *S.b* * | *S.h* | *S.h* | *S.h* |
| J34 | Kachi | Jigawa | Human | *S.h* x *S.b* * | *S.h* | *S.h* | *S.h* |
| J35 | Kachi | Jigawa | Human | *S.h* x *S.b* * | *S.h* | *S.h* | *S.h* |
| J36 | Kachi | Jigawa | Human | *S.h* x *S.b* * | *S.h* | *S.h* | *S.h* |
| J37 | Kachi | Jigawa | Human | *S.h* x *S.b* * | *S.h* | *S.h* | *S.h* |
| J38 | Kachi | Jigawa | Human | *S.h* x *S.b* * | *S.h* | *S.h* | *S.h* |
| J39 | Kachi | Jigawa | Human | *S.h* x *S.b* * | *S.h* | *S.h* | *S.h* |
| J40 | Kachi | Jigawa | Human | *S.h* x *S.b* * | *S.h* | *S.h* | *S.h* |

<sup>1</sup>*S.h* = *S. haematobium*; *S.b* = *S. bovis*; *S.c* = *S. curassoni*; *S.h* x *S.b* = *S. haematobium* x *S. bovis*; *S.h* x *S.c* = *S. haematobium* x *S. curassoni*; \* presence of a private SNP; n.d. = not done
<sup>2</sup>ITS Accession # OQ559623-OQ559638 & OQ564407-OQ564445; Cox1 Accession #OQ568654-OQ568695; ND5 Accession #OQ571658 -OQ571713

In Ile Oluji/Oke Igbo, an endemic site devoid of livestock, all patients were likewise infected with interspecific hybrids. At the *CO1* marker, all isolates possessed alleles that branched with *S. bovis* (Table 2). Phylogenetic analysis at *CO1* showed that infected patients possessed one of 4 unambiguous sequences that branched separately, but clustered tightly with other *S. bovis* sequences, including those from a cow in Nigeria (this study) and from the public domain, all forming a monophyletic group that resolved from other human and livestock schistosomes within the *S. haematobium* species complex with good bootstrap support, including *S. curassoni, S. mattheei, S. intercalatum,* and *S. guineensis* (Figure 2). At *CO1*, one allele was dominant, and the other 3 alleles possessed a single, private SNP indicating that at least 4 haplotypes were circulating among the patient samples (Figure 3). Further, mitonuclear discordance was observed in all 17 isolates, worms possessed an *S. bovis* mitochondrial ancestry, but one of 2 different haplotypes based on the alleles present at the nuclear *transITS* marker, either admixed (heterozygous between *S. haematobium x S. curassoni*) or homozygous for *S. curassoni.* The presence of livestock-specific schistosomes in two patients, the result of hybridization between *S. bovis* x *S. curassoni* with no evidence of *S. haematobium* was noteworthy but has been reported previously in Niger [22]. The other 15 isolates, however, were consistent with livestock:human schistosome hybrids. Our data show that a diverse array of hybrid livestock:human schistosomes are circulating in Nigeria, and they are causing infections in non-pastoral sites, devoid of livestock.

### Bravo-Figey (BF) domain-containing nuclear and ND5 mitochondrial markers

To better understand the true number of haplotypes present, and to further resolve the origin of the *S. bovis* x *S. curassoni* hybrids, we developed two additional genetic markers. The first, *BF*, a cell adhesion molecule (CAM) that belongs to the N-CAM transmembrane proteins, represents an unlinked marker located on chromosome 7. The gene has seven exons, is 4164 nucleotides in length, and PCR primers were designed in a nested configuration to amplify a polymorphic fragment within exon 2 that is 408 nucleotides in length. The genus-specific primers amplify all *Schistosoma* species with equal efficiency (data not shown). DNA sequencing was next used to distinguish between the species, of which 29, 14, 9 and 3 SNPs separate the *S. haematobium* allele from that of *S. mansoni*, *S. bovis*, *S. curassoni* and *S. mattheei* reference sequences, respectively (Figure 4). In Kachi, all isolates but one possessed an unambiguous, homozygous *S. haematobium BF* allele whereas isolate J15 was heterozygous and had an *S. bovis* x *S. haematobium* allele (Table 2). In total, only two hybrid *Sb x Sh* haplotypes were resolved in Kachi among the 4 markers applied (Table 2). In contrast, in Ile Oluji/Oke Igbo, 3 allelic types were resolved at *BF,* either a homozygous *S. haematobium* allele in 6 of 17 isolates, a heterozygous admix of *S. haematobium* x *S. bovis* alleles in 10 of 16 isolates, and one heterozygous admix of *S. haematobium* x *S. curassoni* in the final isolate (Table 2). Importantly, in the two isolates (OD14, OD17) that represented *Sb* x *Sc* livestock:livestock hybrids when assessed at just *CO1* and *ITS* genotyping markers, inclusion of the *BF* marker established that the genotypes for these two isolates were hybrid livestock (*Sb x Sc)* by human (*Sh*) recombinants that had undergone repeated interspecific hybridizations, including at least two separate backcrosses, in the origin of these human infective schistosomes. For the majority of the other Ile Oluji/Oke Igbo isolates, a cross between a *S. haematobium* male with a hybrid *S. bovis x S. curassoni* female would be sufficient to produce the genotype resolved, specifically a maternally inherited *S. bovis* mitochondrial genome that is heterozygous with an *Sh* and either an *Sb* or *Sc* allele at the nuclear markers. Further, in this agrarian community, in the absence of livestock, it may also suggest that people are playing an important role in the origin and evolution of these human infective hybrids. It will be important to genotype individual miracidia isolated from patient samples to determine the extent to which mixed infections are occurring that could promote cross-pairing among the various species within the *S. haematobium* species complex.

**Figure 4.**
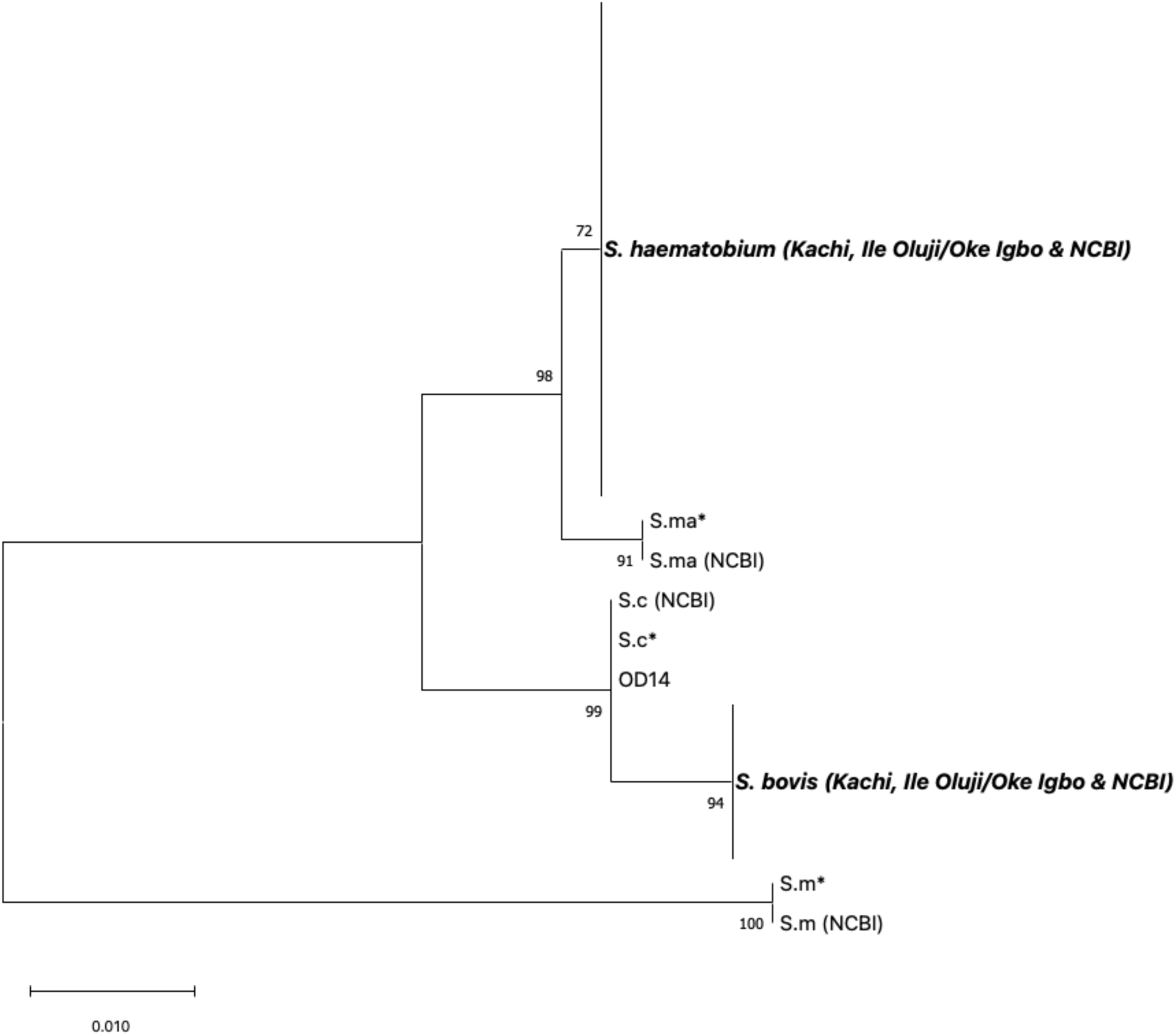
Maximum likelihood tree at BF. Phylogenetic analysis of *BF* alleles recovered from *S. haematobium* isolates in Oluji/Oke Igbo, Ondo state and Kachi, Jigawa state in Nigeria were compared against reference sequences (S.m* = *S. mansoni, S.ma* = S. mattheei, S.c* = S. curassoni* and *S. bovis*). For isolates that were heterozygous, only non-*S.haematobium* alleles are shown in the tree. Hence, for sample OD14, only the *S. curassoni* allele is depicted for the Sh x Sc alleles present at *BF*. Likewise, for OD2,6,8-13, 15-16, and J15, only the *S. bovis* allele is depicted for the Sh x Sb alleles present at *BF*. All other isolates were homozygous for *S. haematobium* alleles at *BF*.

Finally, recombination and rapid *in situ* evolution has been shown to occur in mitochondrial genomes for a variety of parasitic pathogens during interspecific mating [23] or during passage [24]. We developed PCR primers (genus- and *S. bovis*specific primers; Suppl. Fig 3) in a nested configuration to amplify the *ND5* gene within the mitochondrial genome. The genus-specific primers amplified all *Schistosoma* species with equal efficiency (data not shown). When applied against all isolates, no interspecific recombination was resolved, and the *ND5* sequence types were in linkage with the sequence type resolved at *CO1* confirming that the isolates from Kachi had inherited an *S. haematobium* mitochondrial genome, and those from Ile Oluji/Oke Igbo had inherited an *S. bovis* mitochondrial genome (Table 2). Two isolates, one each from Kachi and Ile Oluji/Oke Igbo, each possessed a single private SNP, increasing the number of haplotypes resolved among the patient samples (Suppl. Fig. 2). Ile Oluji/Oke Igbo

## Conclusion

It is apparent that the commonly used CO1 and ITS gene markers are inadequate for correctly characterizing or categorizing *S. haematobium* isolates as either pure or hybrid. Therefore, there is a need to develop more markers and apply them using modern high throughput methods to more precisely determine the extent and number of circulating interspecific hybrids to prioritize these samples for whole genome sequencing. We propose a multilocus typing scheme involving linked and unlinked nuclear gene markers located on the seven *S. haematobium* chromosomes to determine the extent of hybridization that is occurring in schistosomiasis-endemic settings. We also provide compelling evidence that admixed *S. haematobium*:livestock schistosomes are widespread and their origin and transmission may be less dependent on human/cattle contact than previously envisaged. Investigating the genetic diversity and degree of interspecific *S. haematobium* hybrids circulating in snail vector and other animal reservoir hosts (for example, rodents) may provide useful insight in the transmission and origin of these interspecific hybrids. It will also be important to individually genotype individual miracidia derived from infected individuals to assess the degree of mixed infections, and to assess whether cross-pairings with animal:human schistosomes is occurring *in situ* in infected people.

Finally, there is an urgent need to identify a ‘pure’ *S. haematobium* reference sequence (if it exists). It will be important to screen isolates from Zanzibar or Madagascar where no *S. bovis* ancestry has been recorded. This will provide the genomic backbone to compare whole genome sequences of *S. haematobium* populations from different geographical regions, in order to map introgressed and genomic regions that are promoting disease transmission and host range specificity.

## Materials and Methods

### Ethics Statement

This study was conducted under the National ethical permit and protocol numbers NHREC/01/01/2007-30/10/2020 and NHREC/01/01/2007-29/03/2022B. After informed consent by parents and guardians, about 20ml of urine was collected in specimen bottles from a total number of 219 participants ranging in age from four to fourteen years.

### Study sites

Ile Oluji/Oke Igbo is an administrative area of Ondo state situated in the tropical rainforest belt of Southwestern Nigeria. The area which lies between latitude 5° 45N and 8° 15N and longitude 4° 30E is made up of fourteen villages. The climate is humid with small seasonal and daily variations in rainfall. The rainfall is concentrated during the months of May to October with a short break in August and considerable variations from year to year. The area is surrounded by many rivers, such as Owena, Aigo, Esinmu, Iyire, Ogburu, Oni and Awo. The inhabitants of the study area are primarily farmers, engaging in small to medium scale production of both cash and arable crops like yam, cassava, kolanut, cocoa and maize among others [15] (Fig. 1).

Kachi is a village situated in Northern Nigeria under Dutse Local Government Area of Jigawa State (latitude 11° 73’ N and longitude 9° 33’ E). Dutse is a city located in Northern Nigeria and is the capital of Jigawa State. The population is predominantly the Hausa and Fulani tribes practicing agriculture and pastoralism as the main source of livelihood. The climate of the area is tropical wet and dry, and the vegetation type is Sudan Savannah despite its rocky topography typical of Dutse. Over the course of the year, the temperature typically varies reaching highs of 39°C [16] (Fig 1).

### Parasitological analysis

Urine (20ml) was collected in specimen bottles labeled with a unique identifier number on a pre-designed epidemiological form. The sex and age of the participants was entered against the appropriate number on the epidemiological form when urine samples were submitted. In the laboratory, identification of *S. haematobium* eggs was based on their characteristic terminal spine, and both the prevalence and infection intensity were calculated by microscopy. Infection intensity was determined as the number of eggs detected per 10 ml of urine (eggs/10 ml). Light infection was categorized as 1–49 eggs/10 ml and heavy infections ≥50 eggs/10 ml [17]. Urine positive samples for *S. haematobium* eggs were subsequently filtered using Sterlitech 13 mm polycarbonate screen membrane filters (https://www.sterlitech.com/schistosome-test-kit.html) for collection of eggs and preserved in absolute ethanol for molecular analysis.

### Molecular analysis

Total DNA was extracted from individual filters and five *Schistosoma* species were used as positive controls (*S. mansoni*, *S. curassoni*, S. *bovis*, S. *mattheei,* and *S. haematobium*) using QAIGEN Blood and Tissue kit following the manufacturers protocol (www.qiagen.com). The filters were incubated at 56℃ in lysis buffer overnight prior to completing the DNA isolation. Isolated DNA was eluted in molecular grade water and stored at -20℃. Partial region of the mitochondrial cytochrome oxidase subunit 1 (*CO1*) and the complete nuclear ribosomal internal transcribed spacers (trans*ITS*) gene regions was amplified for the isolates using published primers Cox1_schist_5ʹ (5ʹ-TCTTTRGATCATAAGCG-3ʹ), Cox1_schist_3ʹ (5ʹ-TAATGCATMGGAAAAAAACA-3ʹ) and ETTS1 (5’- TGCTTAAGTTCAGCGGGT-3’), ETTS2 (5’-TAACAAGGTTTCCGTAGGTGAA-3’), respectively [18, 19]. We designed new pan-genus primers targeting the mitochondrial *Nad5* gene (5ʹ-GGGTAAAAGTTGGAATTTGAGGG-3ʹ and 5ʹ- CGCTTTAACCATCTGACCACC-3’) and an *S. bovis*-specific (5ʹ-TCGATTTGGAGATGTGGCGT-3ʹ and 5ʹ-ACTGAACTAAA- GCCAAGTCTACC-3ʹ) primer to provide confirmatory data sets for validating Cox1 data sets that do not possess an *S. haematobium* allele. We also identified a new nuclear gene marker, *Bravo_Figey* domain containing protein gene (*BF*) and designed a nested primer set (EXT: 5ʹ-TGTATCACGCTGGCCATACT-3ʹ and 5ʹ-CCAC- CTGCCATCAAACTCAC-3’; INT: 5ʹ-ACTAGATGGCAGATACGGACC-3ʹ and 5ʹ-TAGTCCCCTTGAGGTTGTCG-3’) to amplify the polymorphic region between *S. haematobium* and *S. bovis*. We also sequenced DNA from our five other reference Schistosome species to obtain sequences for phylogenetic analysis. We used the following PCR cycling conditions for the NAD5 and BF primers, 3 minutes at 98°C followed by 35 cycles each of 20 seconds at 98°C, 15 seconds at 58°C followed by 30 seconds at 72°C, and a final elongation step at 72°C for 1 minute. All PCRs were performed in a 25ul reaction using KAPA HIFI Taq reagents, 0.25ul of 50um forward and reverse primers and 1ul of DNA. 4ul of amplicons was visualized on agarose gel by gel electrophoresis. Amplicons were purified using Ampure XP beads and Sanger sequenced in both directions.

### Genetic analysis

All sequences were imported into Geneious vs 2022.2.1 for de novo assembly and trimming. Polymorphic positions in consensus sequences were crosschecked by visualizing the original chromatograms of the forward and reverse sequences. The NCBI nucleotide blast tool (https://blast.ncbi.nlm.nih.gov) was used to initially confirm the identity of each consensus sequence. Several published *Schistosoma* species sequences were retrieved from the NCBI nucleotide database (Suppl. Table 1) for phylogenetic and network analysis with the novel sequence datasets from this study (Table 2). All sequences were aligned using MUSCLE program within the Geneious software prior to subsequent analysis. The molecular phylogenetic placement of the sequence data was implemented in MEGA v11 [20], using Maximum Likelihood method at 1000 bootstrap replicates to ascertain the reliability of the tree branches. Haplotype network was implemented in an R environment using the Pegas package [21].

## Acknowledgements

We would like the thank the Natural History Museum, London for the genetic material provided via the Schistosomiasis Collection at the Natural History Museum (SCAN) archive, it was curated and collected in a collaborative effort, and we acknowledge the support and generosity of the partnerships involved in the collections, particularly in relation to the endemic countries involved, and past colleagues that maintained the live material. The Schistosomiasis Collection at the Natural History Museum (SCAN) was funded by the Wellcome Trust (grant 104958/Z/14/Z). We would also like to thank the Schistosome Resource Centre of the Biomedical Research Institute (sponsored by NIAID, NIH), and in particular, Dr. Margaret Mentink-Kane.

## Conflict of interest

The authors declare that the research was conducted in the absence of any commercial or financial relationships that could be construed as a potential conflict of interest.

## Supporting information

All supporting information can be found in the Supplementary Figures and Tables provided.

## Financial Disclosure Statement

This work was supported by an International Foundation of Science (IFS) grant I3-B-6522-1 and by the Division of Intramural Research within the National Institute of Allergy and Infectious Diseases (NIAID) at the National Institutes of Health (NIH).

## Author contributions

AOG- Conceptualization, Formal Analysis, Data Curation, Visualization, Original Draft Preparation

EEE- Formal Analysis, Original Draft Preparation

BJB- Investigation

OOT- Investigation

GME- Data Curation, Supervision, Validation, Review & Editing, Resources

---

**Supplementary Figure 1.**
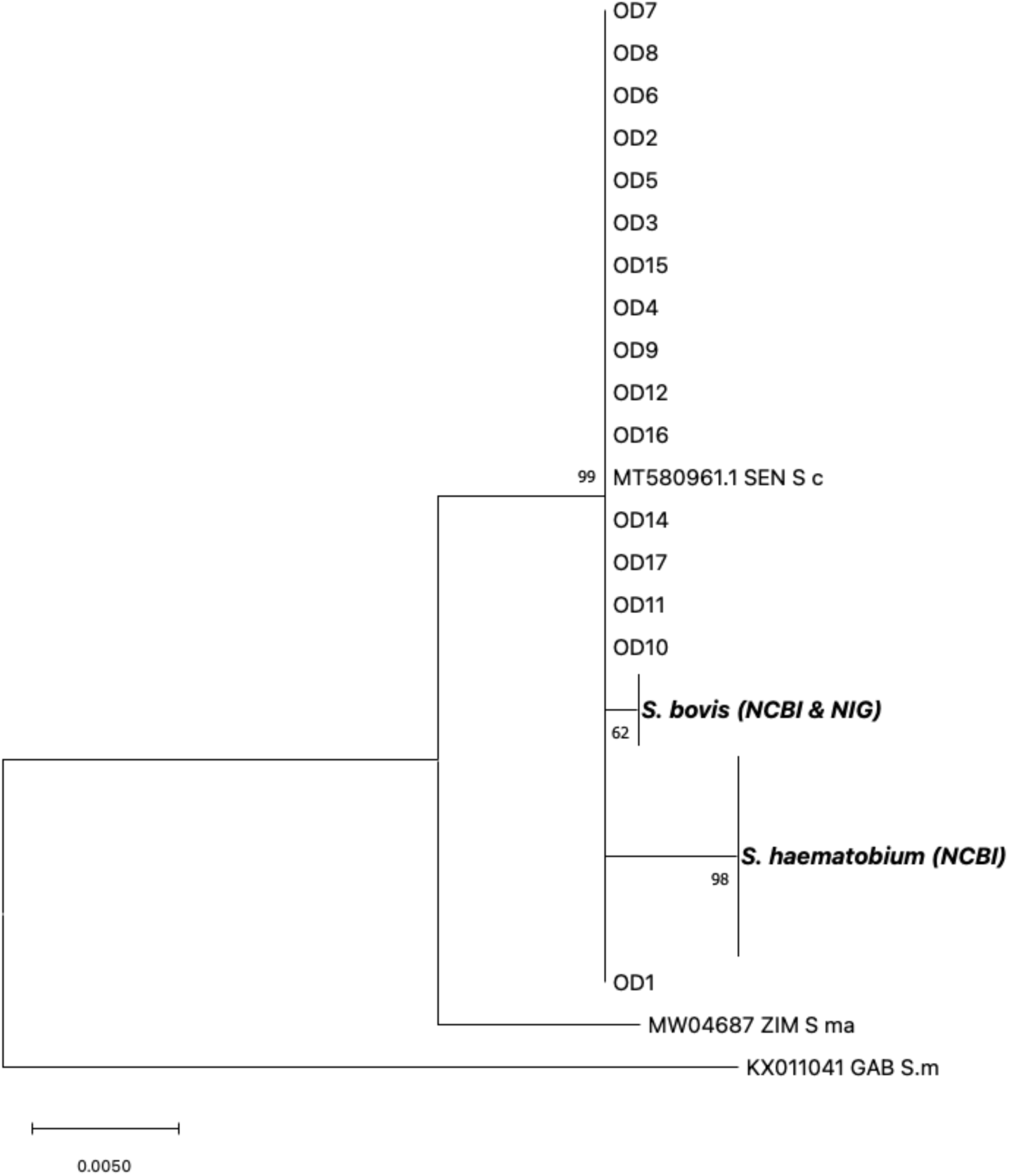
Maximum likelihood tree at the *transITS* marker for *S. haematobium* isolates in Oluji/Oke Igbo (OD). For isolates OD14 and OD17, only an *S. curassoni* allele was recovered at *transITS.* All other isolates were heterozygous and possessed an *S. haematobium* allele x *S. curassoni* allele. Only the *S. curassoni* allele from OD is depicted for comparison against reference alleles for the following: *S. bovis* Nig = *S. bovis* worm recovered from a cow in Nigeria. *S. bovis* (NCBI) = MW027649.1, MT158872.1, MW027648.1, MF776588.1, MF776589.1. *S. haematobium* (NCBI) = MH014047, GU257398, JQ397400, JQ397401, JQ397402, JQ397403, JQ397404, JQ397405, JQ397406, JQ397407, JQ397408, JQ397409, JQ397410, JQ397411, JQ397412, JQ397413, JQ397414. S.c = *S. curassoni* isolate from SEN (Senegal), GenBank Accession number MT580961. S. ma = *S. mattheei* isolate from ZIM (Zimbabwe), GenBank Accession number MW04687. S. m = *S. mansoni* isolate from GAB (Gabon), GenBank Accession number KX011041.

**Suppl. Figure 2:**
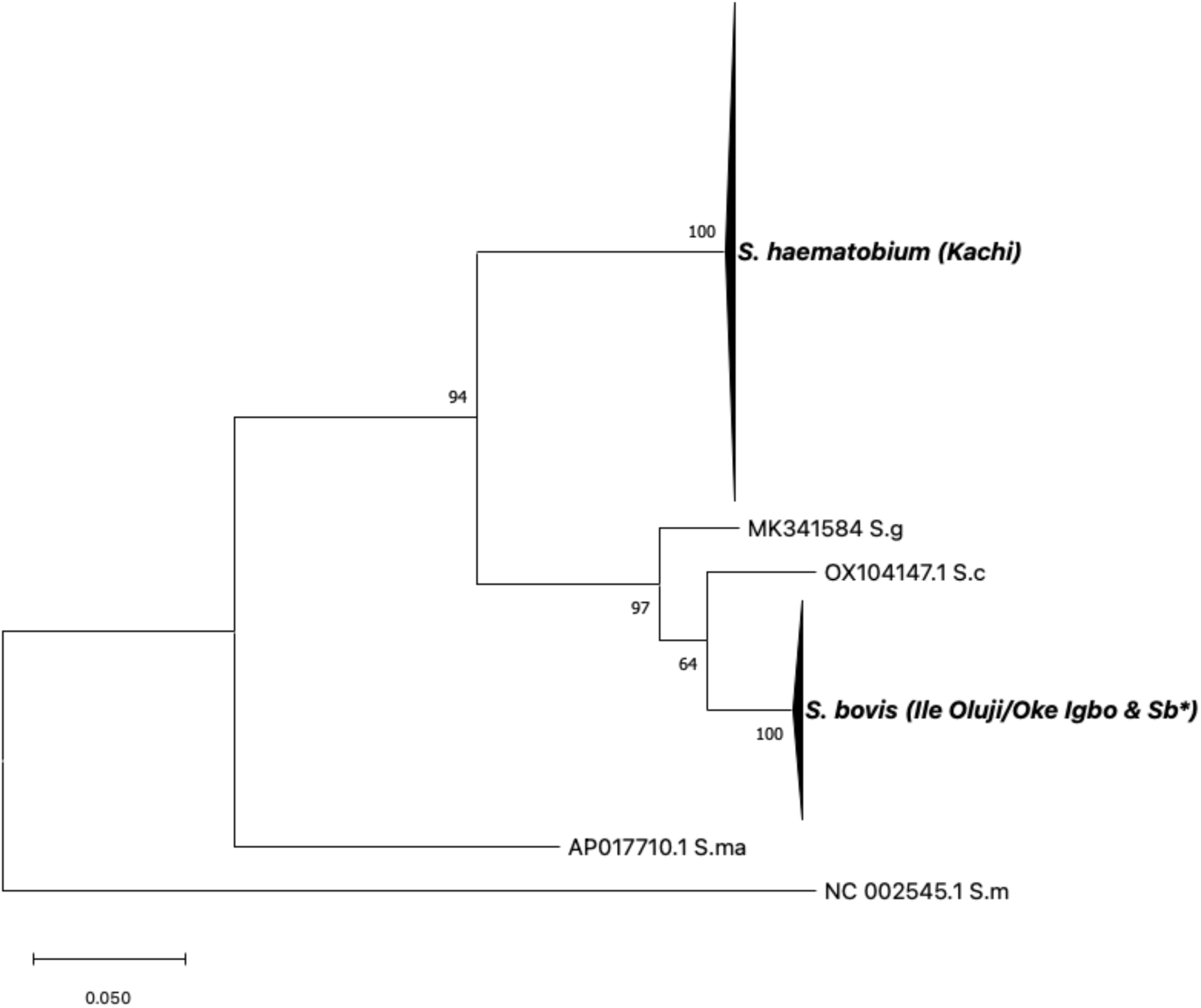
Maximum likelihood tree at ND5. Phylogenetic analysis of *ND5* alleles recovered from *S. haematobium* isolates in Oluji/Oke Igbo (OD) and Kachi, (J) in Nigeria. Only *S. bovis* alleles were identified among isolates recovered from Oluji/Oke Igbo (OD), whereas only *S. haematobium* alleles were identified among isolates recovered from Kachi, (J). * This study, (*S. bovis* worm from cow in Nigeria), S.g = *S. guineensis*, S.c = *S. curassoni*, S.m =*S. mansoni*

**Suppl. Figure 3:**
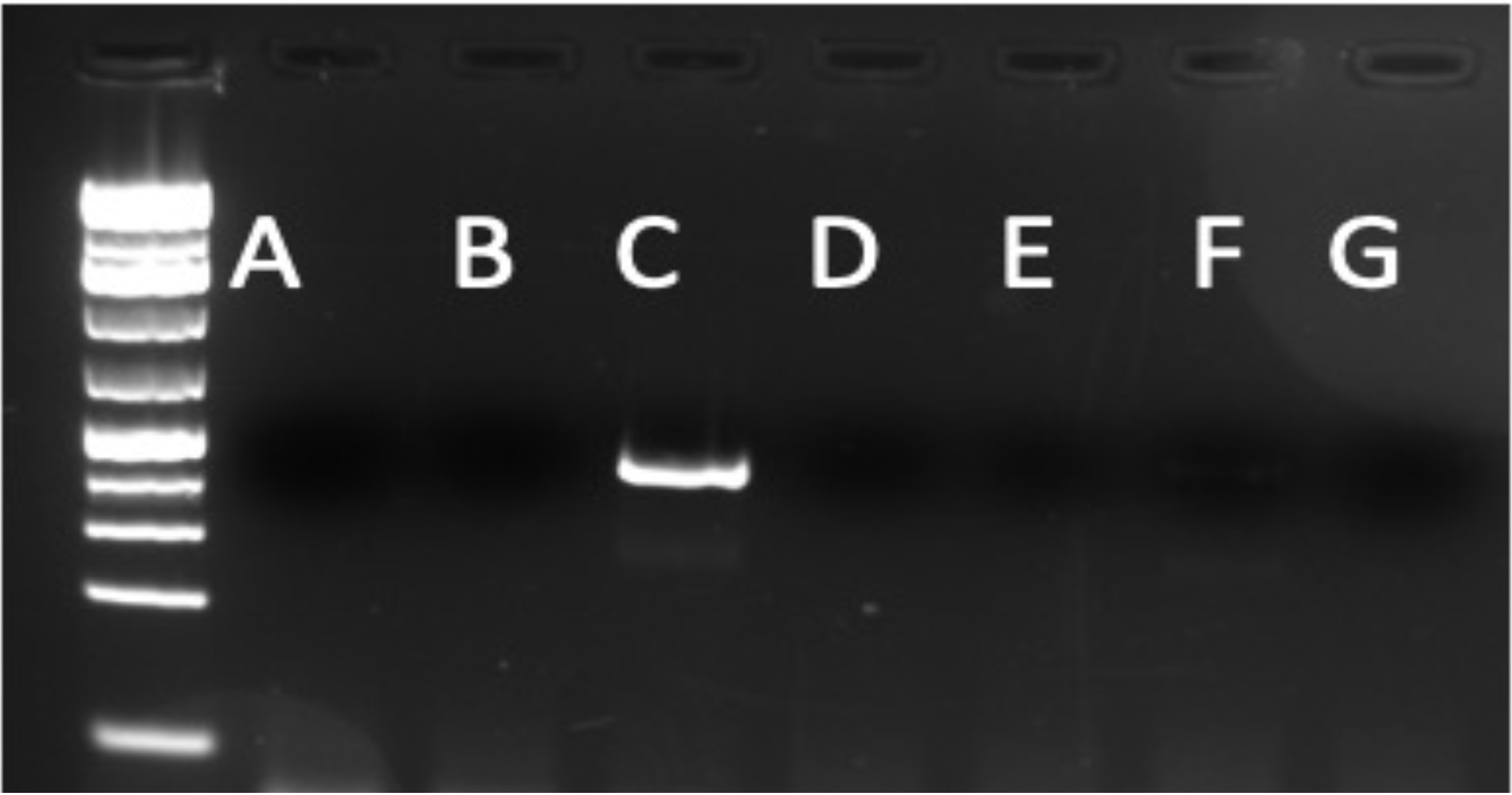
*ND5 S. bovis* specific primer validation. New primers developed at *ND5* were exclusively specific for *S. bovis.* An agarose gel stained with GelRed shows a PCR product only for the PCR reaction that used template DNA from *S. bovis.* Each well used the following template DNA for the PCR reaction: A (*S.haematobium);* B (*S. mansoni*); C (*S. bovis);* D (*S. japonicum);* E (*S. mattheei);* F (*S. curassoni);* G (Water)

**Supplementary Table 1:** CO1 haplotypes based on Fig 3.

| Haplotype | S. h HG 1 | S. h HG 2 | J (Kachi) | OD (Ile Oluji/Oke Igbo) | S. bovis | S. b * (S. bovis , NIG) | S. curassoni | NCBI Sequences in Haplogroups |
| --- | --- | --- | --- | --- | --- | --- | --- | --- |
| I | 10 | 0 | 39 | 0 | 0 | 0 | 0 | GU257337,JQ397332,JQ397345,JQ397350,JQ397351,JQ397352,JQ397353,JQ397397,JQ397369,JQ397392 |
| II | 1 | 0 | 0 | 0 | 0 | 0 | 0 | JQ397349 |
| III | 1 | 0 | 0 | 0 | 0 | 0 | 0 | JQ397368 |
| IV | 0 | 0 | 1 | 0 | 0 | 0 | 0 |  |
| V | 1 | 0 | 0 | 0 | 0 | 0 | 0 | JQ082121 |
| VI | 1 | 0 | 0 | 0 | 0 | 0 | 0 | JQ397393 |
| VII | 0 | 1 | 0 | 0 | 0 | 0 | 0 | JQ397399 |
| VIII | 0 | 5 | 0 | 0 | 0 | 0 | 0 | GU257334,GU257338,GU257348,GU257360,JQ397398 |
| IX | 0 | 1 | 0 | 0 | 0 | 0 | 0 | GU257346 |
| X | 0 | 1 | 0 | 0 | 0 | 0 | 0 | GU257336 |
| XI | 0 | 1 | 0 | 0 | 0 | 0 | 0 | GU257335 |
| XII | 0 | 1 | 0 | 0 | 0 | 0 | 0 | GU257345 |
| XIII | 0 | 1 | 0 | 0 | 0 | 0 | 0 | GU257352 |
| XIV | 0 | 1 | 0 | 0 | 0 | 0 | 0 | GU257354 |
| XV | 0 | 1 | 0 | 0 | 0 | 0 | 0 | GU257356 |
| XVI | 0 | 1 | 0 | 0 | 0 | 0 | 0 | GU257358 |
| XVII | 0 | 1 | 0 | 0 | 0 | 0 | 0 | GU257342 |
| XVIII | 0 | 1 | 0 | 0 | 0 | 0 | 0 | GU257349 |
| XIX | 0 | 0 | 0 | 0 | 0 | 0 | 1 | AJ519516 |
| XX | 0 | 0 | 0 | 0 | 1 | 0 | 0 | MW022138 |
| XXI | 0 | 0 | 0 | 1 | 0 | 0 | 0 |  |
| XXII | 0 | 0 | 0 | 0 | 1 | 0 | 0 | AJ519521 |
| XXIII | 0 | 0 | 0 | 0 | 1 | 0 | 0 | MT579448 |
| XXIV | 0 | 0 | 0 | 11 | 0 | 0 | 0 |  |
| XXV | 0 | 0 | 0 | 0 | 1 | 0 | 0 | MF919415 |
| XXVI | 0 | 0 | 0 | 0 | 1 | 0 | 0 | MK757170 |
| XXVII | 0 | 0 | 0 | 1 | 0 | 0 | 0 |  |
| XXVIII | 0 | 0 | 0 | 1 | 0 | 0 | 0 |  |
| XXIX | 0 | 0 | 0 | 0 | 0 | 1 | 0 |  |
| XXX | 0 | 0 | 0 | 0 | 1 | 0 | 0 | AY157212 |
| XXXI | 0 | 0 | 0 | 0 | 1 | 0 | 0 | FJ897160 |

**Supplementary Table 2:**
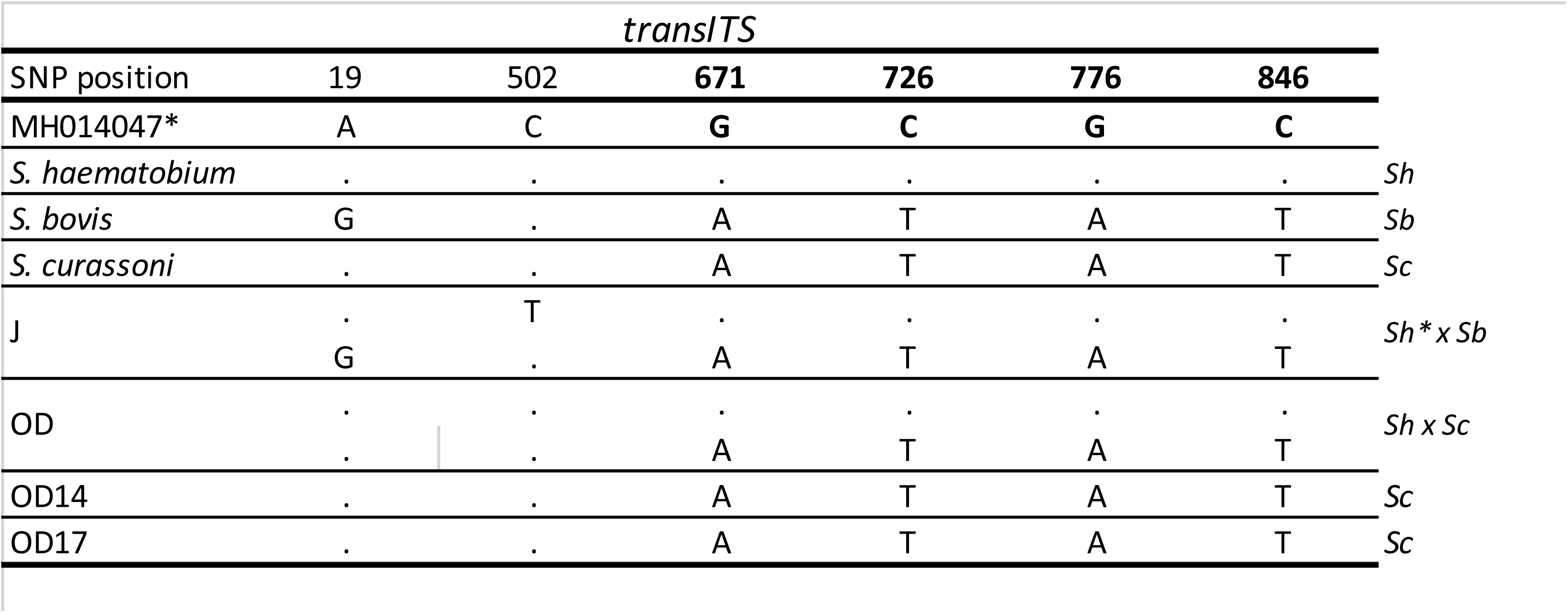
*transITS* haplotypes. * SNP position reference, *S.haematobium* NCBI (MH014047), J = Kachi Isolates, OD = Ile Oluji/Oke Igbo isolates, *S.h* = *S. haematobium*, *S.b* = *S. bovis*.

Ajakaye et al\_Suppl. Table 1
| transITS |  |  |  |  |  |  |  |
| --- | --- | --- | --- | --- | --- | --- | --- |
| SNP position | 19 | 502 | 671 | 726 | 776 | 846 |  |
| MH014047* | A | C | G | C | G | C |  |
| *S. haematobium* | . | . | . | . | . | . | *Sh* |
| *S. bovis* | G | . | A | T | A | T | *Sb* |
| *S. curassoni* | . | . | A | T | A | T | *Sc* |
| J | . | T | . | . | . | . | *Sh\* x Sb* |
|  | G | . | A | T | A | T |  |
| OD | . | . | . | . | . | . | *Sh x Sc* |
|  | . | . | A | T | A | T |  |
| OD14 | . | . | A | T | A | T | *Sc* |
| OD17 | . | . | A | T | A | T | *Sc* |

## Notes

### Competing Interest Statement

The authors have declared no competing interest.

